# Data Processing in Multidimensional MRI For Biomarker Identification: Is It Necessary?

**DOI:** 10.1101/2025.03.25.645236

**Authors:** Kristofor Pas, Dan Benjamini, Peter Basser, Gustavo Rohde

**Affiliations:** Department of Biomedical Engineering, University of Virginia; Multiscale Imaging and Integrative Biophysics Unit, National Institute on Aging, NIH; Section on Quantitative Imaging and Tissue Sciences, National Institutes of Health, NIH

**Keywords:** Machine Learning, Multidimensional MRI, Microstructure, Diffusion

## Abstract

Multidimensional MRI (MD-MRI) is an emerging technique that holds promise for identifying tissue characteristics that could be indicative of pathologies. Before these characteristics can be interpreted, MD-MRI measurements are converted into an spectrum. These spectra are then utilized to obtain some understanding of the underlying tissue microstructure, often through the use of statistical, machine learning, and mathematical modeling methods. The aim of this study was to compare outcomes of using unprocessed MDMRI signals for statistical regression in comparison to the corresponding spectra. Backed by a theoretical argument, we described an experimental procedure regressing both MDMRI signals and spectra to histological outcomes intrasubject. Through using multiple conventional ML methods, and a proposed method using convex sets, we aimed to see which yielded the highest accuracy. Both theory and experimental evidence suggest that, without *a priori* information, statistical regression was best performed on the MDMRI signal. We conclude, barring any *a priori* information regarding tissue changes, there is no significant advantage to performing regression analysis on reconstructed spectra in the process of biomarker identification.

## Introduction

Diffusion–relaxation multidimensional MRI (MD-MRI) combines sensitivity to meso- and microstructural properties(1) with the ability to probe chemical composition.(2) This technique works through varying acquisition parameters to jointly probe correlations among local relaxation and diffusion mechanisms within each voxel. Due to the numerous acquisitions built into the protocol, the signal acquired from MD-MRI can be used to estimate high-dimensional diffusion-relaxation distribution as opposed to a scalar output as in conventional MRI (3) .These dimensions contain data reflecting microstructural and compositional features of different water pools within a given image voxel. The MD-MRI signal can be converted into an probability density function (referred to as a spectrum) that can often be interpreted as a biologically-related component or related to a physiological process.

Because of the data-rich, interpretable output from MD-MRI, it is of interest to use this information to develop quantitative imaging biomarkers. Using MD-MRI spectra, machine learning and statistical methods can parse out nuanced differences at a sub-voxel scale.(4–7) To date, studies have included MD-MRI to find important biomarkers underlying diseased and normative tissue specimen.(8–16) With the biophysically interpretable spectra, it is possible to improve scientific understanding underlying tissue changes. Therefore, it has potential in improving diagnostic methods in both preclinical(17, 18) and clinical(19–21) settings.

However, MD-MRI is not without its pitfalls. The spectra derived from the signal output provides insight into changes in tissue water pools, though comes at a steep price of computation and without certainty of results(22). This inverse problem, based on a Fredholm integral equation of the first kind, is ill-posed, and requires optimization to arrive at an approximate a solution.(23) Experimental design features and experimental parameters, such as low SNR, can all affect our solution. Furthermore, even in instances where information is maintained through numerical inversion of the signal, the classification of distributions is not a well-understood problem in that the data geometry is highly non-linear.

In this methodological note, we aim to elucidate the fact that, barring specific *a priori* knowledge of how diffusion-relaxation distributions may differ between different tissues and states (i.e., normal vs. pathological), the MD-MRI signals are just as informative as the estimated spectra for regression. We demonstrate this experimentally in the specific application of correlating MD-MRI measurements to labels obtained from registered histological sections, though it is our expectation that the findings may apply more generally.

### Theory

In this section, we use established statistical theory to elucidate the limitations of MD-MRI for regressing biomarkers.

#### A. Conventional MDMRI Inversion

Multidimensional MRI (MD-MRI) combines NMR relaxation and diffusion data acquisition with MR-based spatial localization. In *T*_1_− *T*_2_− *D* MRI, often MR volumes are acquired with different longitudinal relaxation, transverse relaxation, and diffusion-weighting parameters, namely inversion time (*τ*_*i*_), echo time (*τ*_*E*_), and b-value (**b**), respectively. These acquisitions create a tuple *β*_*i*_ := (*τ*_*i*_, *τ*_*E*_, **b**)_*i*_ for the *i*-th acquisition. When the *j*-th voxel is samples *d* times, it yields a vector of MD-MRI measurements, 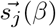

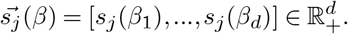

The multidimensional distribution of underlying diffusion-relaxation properties is related to the signal according to the Fredholm Integral Equation of the first kind

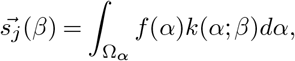

where *f* (*α*) is the joint probability distribution *p*(*T*_1_, *T*_2_, *D*), embodying correlations among the three MR parameters which is weighted by kernel *k*(*α*; *β*). The solution can be approximated by

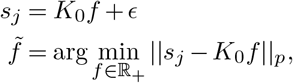

for which || · ||_*p*_ is the p-norm, 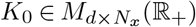 is the kernel matrix, *f*, 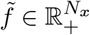 and *ε* ∈ ℝ is a noise process. This problem is ill-posed, solutions are non-unique and sensitive to noise. Therefore, conditioning of the problem through regularization is commonly used with computation (24). Consequently, this is computationally expensive, and depreciates the validity and interpretability of the solutions.

### B. No Clever Manipulations on Data Improves Inference

Let *X* be a random vector associated with a label obtained from pixel intensities in a histological image. Likewise, let *Y* be a random vector corresponding to the MD-MRI signal data. Finally, let *Y* ^/^ correspond to a random vector associated with the distribution data (computationally obtained from the MD-MRI data Y). Our goal is to predict *X* using *Y* or *Y* ^/^. Specifically, we generate predictors 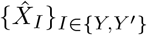 that recapitulate X. Empirical loss 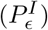 quantifies the predictor quality.

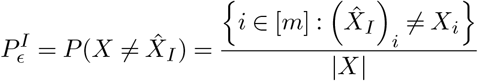

#### Theorem 1

(Fano’s Inequality(25)) Let *X*→*Y* be a function. Suppose 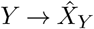 is a predictor for *X* based on *Y*.

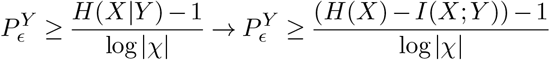

For which *H*(*X*) is the entropy of *X, H*(*X Y*) is the conditional entropy of *X* given *Y, I*(*X*; *Y*) is the mutual information of *X, Y*, and *χ* is the support of *X*.

#### Theorem 2

(Hellman-Raviv’s Theorem (26)) Let *X*→*Y* be a function. Suppose 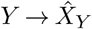 is a Maximum Likelihood Predictor for *X* based on *Y*.

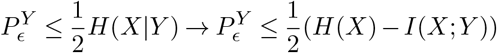

#### Theorem 3

(Data Processing Inequality (25)) Let *X*→ *Y*→*Y* ^/^ be functions.

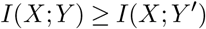

Through direct application of Theorems 1 and 3, we can see that the lower bounds on predictors 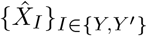 are as follows.

The direct observation from this is that, if the function governing *Y*→ *Y* ^/^ is corrupted, the best possible valid prediction comes from *Y*.

A stronger observation exists in the instance where we have a Maximum Likelihood Predictor, in which we can invoke Theorems 2 and 3 to show the predictions using *Y* can be no worse then predictions based on *Y* ^/^.

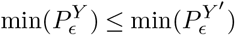

In context of the Maximum Likelihood Predictor, MD-MRI signal 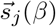 provides less error than the processed spectra (i.e., the inverted signal), 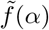 and at worst, provides similar error rates.

### C. A Simple, Useful Algorithm to Classify Data of Different Types

Let *D* = {*µ, g*(*µ*)} ⊂ ℝ^*d*^×𝒞 be a set containing a data distribution, *µ* and a corresponding label function, *g*(*µ*).

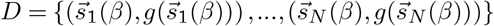

In conventional Regression/Machine Learning, we use an algorithm, *A*, that learns information from *D*. Specifically, *A* seeks to solve for some hypothesis, *h*^∗^ that minimizes loss

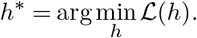

In context of empirical error, we have the following

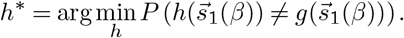

However, finding a decision boundary is intractable, requiring ample training time and data. Furthermore, decision boundaries may be inconsistent between runs, depending on parameters. All of these problems are exacerbated when the data geometry is complex.

To circumvent these problems, we describe a K-Nearest Local Convex Set algorithm (27), which can classify non-linear data geometries without conventional training.

Let *x*^∗^ be a voxel of interest. Within our training set, we have a set of labels corresponding {*C*_*i*_}_*i*∈[*M*]_ to a known distribution. Define 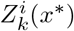 as the K-Nearest neighborhood to *x*^∗^ corresponding to only points in class *C*_*i*_ in our known data. This allows for the computation from *x*^∗^ to each K-Nearest Local Convex set. We establish the hypothesis for this classifier, *h*, as follows

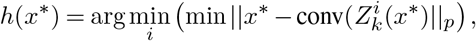

for which conv(·) implies the convex hull of the data.

**Fig. 1.**
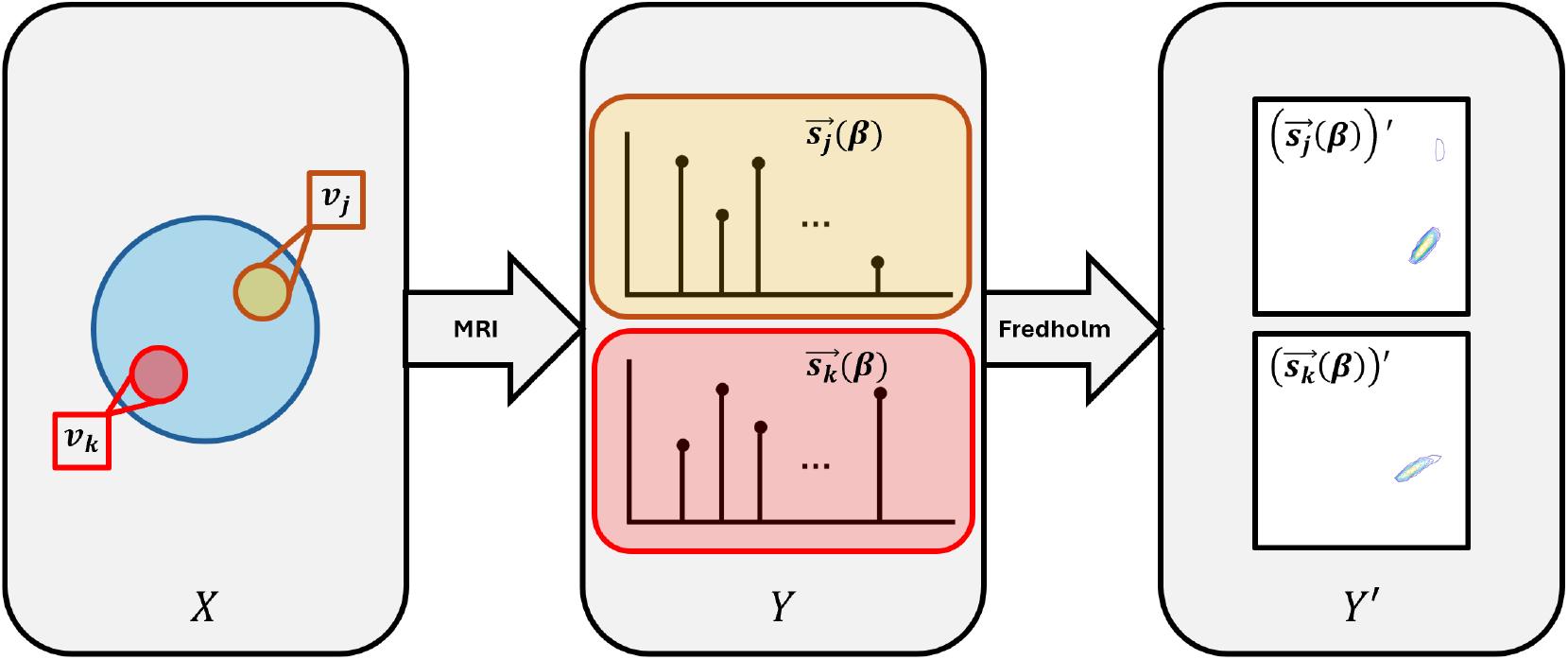
Overview of Conventional Processing within the Field

**Fig. 2.**
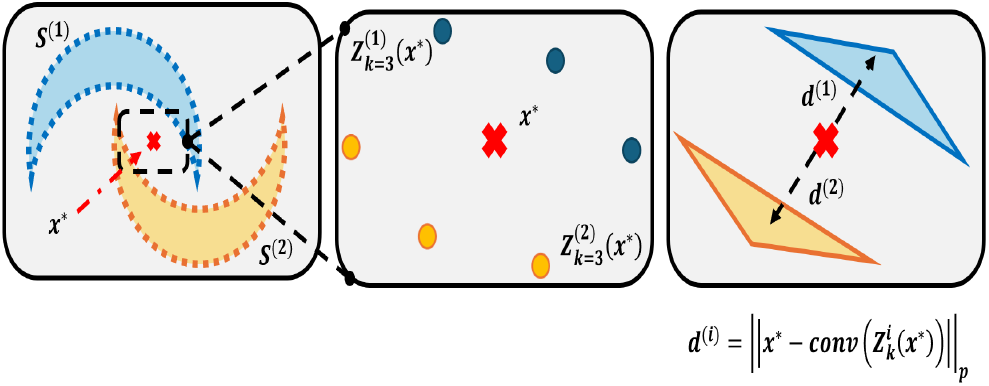
Illustration of Nearest Local Convex Set Algorithm.

**Fig. 3.**
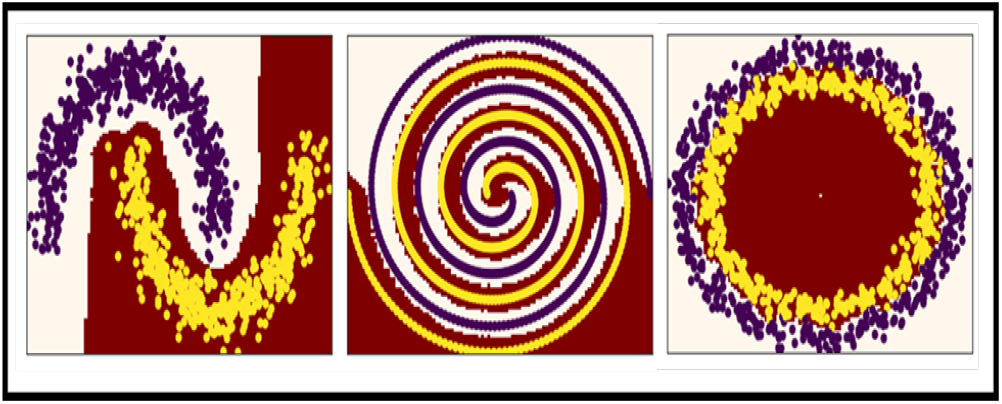
Example NLCS Classification Boundaries with Complex Geometries.

## Methods

### D. Experimentation Protocol

The MRI and microscopy data used in this study were originally reported in Benjamini *et al*.(28) In summary, we analyzed formalin-fixed brain tissue samples from 14 autopsy donors. All experiments were conducted in compliance with Institutional Review Board (IRB) guidelines prior to study initiation. Among the 14 cases, 7 exhibited interface astrogliosis based on prior neuropathological assessment, while the remaining 7 served as controls with no evidence of interface astrogliosis. We conducted a combined postmortem MD-MRI and histopathology study, acquiring MRI data using a 7T Bruker vertical bore scanner. Following MRI acquisition, each tissue block underwent histopathological processing. MD-MRI data were collected using a previously established sampling protocol.(7, 29, 30) Immunohistochemistry for glial fibrillary acidic protein (GFAP) was performed using a Leica Bond III automated immunostainer with a diaminobenzidine chromogen detection system (DS9800, Leica Biosystems, Buffalo Grove, IL) to assess the presence of astrogliosis. Microscopy and MRI images were then co-registered for further analysis. (28)

### E. Methods Related to Classification

For each method, n=30 bootstrapped samples were acquired from N=14 different subjects. Data was then split into 50/50 training/testing instances for each sample. Regression was then performed with each different model. Random States were iteratively defined across different samples to assure consistency of comparing spectra, allowing for paired comparison.

## Results

To test these theories, we investigated the task of regressing MD-MRI voxel values to astrogliosis-related histology pixel values. That is, we utilized statistical regression (i.e., machine learning) models to recover a function from both the raw MD-MRI signal data, Y, and processed (distribution) data, Y’ to the pixel value from the histology data X. For this task, we utilized a dataset from a previous study examining MD-MRI signatures of astrogliosis in human ex vivo subjects (see Benjamini *et al*.(28) for details).

While the maximum likelihood in theory is an optimal framework to produce such predictors, we note that the formulas for the likelihood functions for each *X, Y, andY* ^/^ are not known. To that extent we utilize a series of machine learning models, including a nearest local convex set one described above, to assess whether there is any advantage (or disadvantage) of performing regression based on the raw MD-MRI data (*Y*) over the processed spectral data (*Y* ^/^). To do this, we compared the use of each of the commonly used 2D correlation distributions, namely *T*_1_ −*T*_2_, *T*_1_− *D, T*_2_ −*D*. The score for determining regression quality was Cohen’s Kappa Statistic.

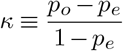

## Discussion

Diffusion-relaxation MD-MRI provides information of tissue microstructure. Previously, it has been shown that using spectra can be regressed to histology.(10, 11, 28, 30) But a more practical method exists without sacrificing, and potentially improving, inference from data. We have shown, with both theory and computational experiments, that in the absence of *a priori* knowledge, regression problems could use unprocessed data representations without loss of accuracy, and potential improvements. Furthermore, we have shown there exist multiple good candidates that can regress this type of data.

**Table 1.** Average Cohen’s Kappa Statistic for Different Machine Learning Techniques on Varying Data Representations. * Indicates Column-wise maximum, **BOLD** Indicates Row-Wise maximum.

| Technique | T1-D | T2-D | T1-T2 | Proposed |
| --- | --- | --- | --- | --- |
| 2-NN | 0.432 | 0.363 | 0.432 | **0.502** |
| MLP | 0.418 | 0.407 | 0.462 | **0.522** |
| LDA | 0.204 | 0.289 | 0.241 | **0.549** |
| NLCS | **0.537\*** | **0.469\*** | **0.517\*** | **0.601\*** |

The current designed study was performed as a proof-of-concept methods for testing the theory described. In its current state, we have shown evidence supporting using unprocessed data, rather than spectral data, for biomarker identification within individual subjects. Further potential studies could iterate on similar ideas, with the intention of studying cross-patient biomarkers.

## Bibliography

1. Paul T. Callaghan. Translational Dynamics and Magnetic Resonance. Oxford University Press, 9 2011. ISBN 9780199556984. doi: 10.1093/acprof:oso/9780199556984.001.0001.

2. P Tofts. Quantitative MRI of the Brain. Wiley, 8 2003. ISBN 9780470847213. doi: 10.1002/0470869526.

3. D. Benjamini and P. J. Basser. Multidimensional correlation mri. NMR in Biomedicine, 33, 2020. ISSN 10991492. doi: 10.1002/nbm.4226.

4. Randall M. Kroeker and Mark R. Henkelman. Analysis of biological nmr relaxation data with continuous distributions of relaxation times. Journal of Magnetic Resonance (1969), 69: 218–235, 1986. ISSN 00222364. doi: 10.1016/0022-2364(86)90074-0.

5. S Peled, D G Cory, S A Raymond, D A Kirschner, and F A Jolesz. Water diffusion, t(2), and compartmentation in frog sciatic nerve. Magnetic Resonance in Medicine, 42:911–8, 11 1999. ISSN 0740-3194.

6. Matthew D. Silva, Karl G. Helmer, Jing-Huei Lee, Sam S. Han, Charles S. Springer, and Christopher H. Sotak. Deconvolution of compartmental water diffusion coefficients in yeast-cell suspensions using combined t1 and diffusion measurements. Journal of Magnetic Resonance, 156:52–63, 5 2002. ISSN 10907807. doi: 10.1006/jmre.2002.2527.

7. Kristofor Pas, Michal E. Komlosh, Daniel P. Perl, Peter J. Basser, and Dan Benjamini. Retaining information from multidimensional correlation mri using a spectral regions of interest generator. Scientific Reports, 10:3246, 12 2020. ISSN 2045-2322. doi: 10.1038/s41598-020-60092-5.

8. Daeun Kim, Eamon K. Doyle, Jessica L. Wisnowski, Joong Hee Kim, and Justin P. Haldar. Diffusion-relaxation correlation spectroscopic imaging: A multidimensional approach for probing microstructure. Magnetic Resonance in Medicine, 78:2236–2249, 12 2017. ISSN 07403194. doi: 10.1002/mrm.26629.

9. Dan Benjamini and Peter J. Basser. Magnetic resonance microdynamic imaging reveals distinct tissue microenvironments. NeuroImage, 163:183–196, 12 2017. ISSN 10538119. doi: 10.1016/j.neuroimage.2017.09.033.

10. Dan Benjamini, Elizabeth B. Hutchinson, Michal E. Komlosh, Courtney J. Comrie, Susan C. Schwerin, Guofeng Zhang, Carlo Pierpaoli, and Peter J. Basser. Direct and specific assessment of axonal injury and spinal cord microenvironments using diffusion correlation imaging. NeuroImage, 221:117195, 11 2020. ISSN 10538119. doi: 10.1016/j.neuroimage.2020.117195.

11. Dan Benjamini, Diego Iacono, Michal E Komlosh, Daniel P Perl, David L Brody, and Peter J Basser. Diffuse axonal injury has a characteristic multidimensional mri signature in the human brain. Brain, 144:800–816, 4 2021. ISSN 0006-8950. doi: 10.1093/brain/awaa447.

12. Alexis Reymbaut, Jeffrey Critchley, Giuliana Durighel, Tim Sprenger, Michael Sughrue, Karin Bryskhe, and Daniel Topgaard. Toward nonparametric diffusion-characterization of crossing fibers in the human brain. Magnetic Resonance in Medicine, 85:2815–2827, 5 2021. ISSN 0740-3194. doi: 10.1002/mrm.28604.

13. Paddy J. Slator, Marco Palombo, Karla L. Miller, Carl-Fredrik Westin, Fredrik Laun, Daeun Kim, Justin P. Haldar, Dan Benjamini, Gregory Lemberskiy, João P. de Almeida Martins, and Jana Hutter. Combined diffusion-relaxometry microstructure imaging: Current status and future prospects. Magnetic Resonance in Medicine, 2021. doi: 10.1002/mrm.28963.

14. Xiaobin Wei, Li Zhu, Yanyan Zeng, Ke Xue, Yongming Dai, Jianrong Xu, Guiqin Liu, Fang Liu, Wei Xue, Dongmei Wu, and Guangyu Wu. Detection of prostate cancer using diffusion-relaxation correlation spectrum imaging with support vector machine model – a feasibility study. Cancer Imaging, 22:77, 12 2022. ISSN 1470-7330. doi: 10.1186/s40644-022-00516-9.

15. Paddy J. Slator, Daniel Cromb, Laurence H. Jackson, Alison Ho, Serena J. Counsell, Lisa Story, Lucy C. Chappell, Mary Rutherford, Joseph V. Hajnal, Jana Hutter, and Daniel C. Alexander. Non-invasive mapping of human placenta microenvironments throughout pregnancy with diffusion-relaxation mri. Placenta, 144:29–37, 12 2023. ISSN 01434004. doi: 10.1016/j.placenta.2023.11.002.

16. Shinjini Kundu, Stephanie Barsoum, Jeanelle Ariza, Amber L Nolan, Caitlin S Latimer, C Dirk Keene, Peter J Basser, and Dan Benjamini. Mapping the individual human cortex using multidimensional mri and unsupervised learning. Brain Communications, 5, 11 2023. ISSN 2632-1297. doi: 10.1093/braincomms/fcad258.

17. Maxime Yon, João P. de Almeida Martins, Qingjia Bao, Matthew D. Budde, Lucio Frydman, and Daniel Topgaard. Diffusion tensor distribution imaging of an in vivo mouse brain at ultrahigh magnetic field by spatiotemporal encoding. NMR in Biomedicine, 33, 11 2020. ISSN 0952-3480. doi: 10.1002/nbm.4355.

18. Maxime Yon, Omar Narvaez, Daniel Topgaard, and Alejandra Sierra. In vivo rat brain mapping of multiple gray matter water populations using nonparametric <b>d</b> (<i>ω</i>)-<i>r</i> <sub>1</sub> - <i>r</i> <sub>2</sub> distributions mri. NMR in Biomedicine, 38, 1 2025. ISSN 0952-3480. doi: 10.1002/nbm.5286.

19. Jan Martin, Alexis Reymbaut, Manuel Schmidt, Arnd Doerfler, Michael Uder, Frederik Bernd Laun, and Daniel Topgaard. Nonparametric d-r1-r2 distribution mri of the living human brain. NeuroImage, 245:118753, 12 2021. ISSN 10538119. doi: 10.1016/j.neuroimage.2021.118753.

20. Jessica T. E. Johnson, M. Okan Irfanoglu, Eppu Manninen, Thomas J. Ross, Yihong Yang, Frederik B. Laun, Jan Martin, Daniel Topgaard, and Dan Benjamini. In vivo disentanglement of diffusion frequency-dependence, tensor shape, and relaxation using multidimensional <scp>mri</scp>. Human Brain Mapping, 45, 5 2024. ISSN 1065-9471. doi: 10.1002/hbm.26697.

21. Eppu Manninen, Shunxing Bao, Bennett A. Landman, Yihong Yang, Daniel Topgaard, and Dan Benjamini. Variability of multidimensional diffusion–relaxation mri estimates in the human brain. Imaging Neuroscience, 2:1–24, 12 2024. ISSN 2837-6056. doi: 10.1162/imag_a_00387.

22. Di Yuan and Xinming Zhang. An overview of numerical methods for the first kind fredholm integral equation. SN Applied Sciences, 1:1–12, 2019.

23. Dan Benjamini. Nonparametric inversion of relaxation and diffusion correlation data, 2020.

24. Abdul-Majid Wazwaz. The regularization method for fredholm integral equations of the first kind. Computers & Mathematics with Applications, 61(10):2981–2986, 2011.

25. Thomas M Cover. Elements of information theory. John Wiley & Sons, 1999.

26. Martin Hellman and Josef Raviv. Probability of error, equivocation, and the chernoff bound. IEEE Transactions on Information Theory, 16(4):368–372, 1970.

27. Pascal Vincent and Yoshua Bengio. K-local hyperplane and convex distance nearest neighbor algorithms. Advances in neural information processing systems, 14, 2001.

28. Dan Benjamini, David S Priemer, Daniel P Perl, David L Brody, and Peter J Basser. Mapping astrogliosis in the individual human brain using multidimensional mri. Brain, 8 2022. ISSN 0006-8950. doi: 10.1093/brain/awac298.

29. Dan Benjamini and PJ Basser. Use of marginal distributions constrained optimization (madco) for accelerated 2d mri relaxometry and diffusometry. Journal of Magnetic Resonance, 271:40–45, 2016. ISSN 10960856 10907807. doi: 10.1016/j.jmr.2016.08.004.

30. Dan Benjamini, Mustapha Bouhrara, Michal E. Komlosh, Diego Iacono, Daniel P. Perl, David L. Brody, and Peter J. Basser. Multidimensional mri for characterization of subtle axonal injury accelerated using an adaptive nonlocal multispectral filter. Frontiers in Physics, 9, 9 2021. ISSN 2296-424X. doi: 10.3389/fphy.2021.737374.

